# Tracktor: image-based automated tracking of animal movement and behaviour

**DOI:** 10.1101/412262

**Authors:** Vivek Hari Sridhar, Dominique G. Roche, Simon Gingins

**Affiliations:** Department of Collective Behaviour, Max Planck Institute for Ornithology, Konstanz, Germany; Department of Biology, University of Konstanz, Konstanz, Germany; Institute of Biology, University of Neuchâtel, 2000 Neuchâtel, Switzerland

**Keywords:** collective behaviour, fast start escape response, locomotion, kinematics, choice experiment, video analysis software

## Abstract

1. Automated movement tracking is essential for high-throughput quantitative analyses of the behaviour and kinematics of organisms. Automated tracking also improves replicability by avoiding observer biases and allowing reproducible workflows. However, few automated tracking programs exist that are open access, open source, and capable of tracking unmarked organisms in noisy environments.

2. *Tracktor* is an image-based tracking freeware designed to perform single-object tracking in noisy environments, or multi-object tracking in uniform environments while maintaining individual identities. *Tracktor* is code-based but requires no coding skills other than the user being able to specify tracking parameters in a designated location, much like in a graphical user interface (GUI). The installation and use of the software is fully detailed in a user manual.

3. Through four examples of common tracking problems, we show that Tracktor is able to track a variety of animals in diverse conditions. The main strengths of Tracktor lie in its ability to track single individuals under noisy conditions (e.g. when the object shape is distorted), its robustness to perturbations (e.g. changes in lighting conditions during the experiment), and its capacity to track multiple individuals while maintaining their identities. Additionally, summary statistics and plots allow measuring and visualizing common metrics used in the analysis of animal movement (e.g. cumulative distance, speed, acceleration, activity, time spent in specific areas, distance to neighbour, etc.).

4. *Tracktor* is a versatile, reliable, easy-to-use automated tracking software that is compatible with all operating systems and provides many features not available in other existing freeware. Access *Tracktor* and the complete user manual here: https://github.com/vivekhsridhar/tracktor

## 1. Introduction

Quantifying animal movement is central to answering numerous research questions in ethology, behavioural ecology, ecophysiology, biomechanics, neuroscience and evolutionary ecology. Movement is also increasingly used as a proxy to evaluate sub-lethal effects of pharmaceuticals, climate change and other anthropogenic stressors on animals, thereby improving our understanding of contemporary environmental and health issues of global importance (Wong & Candolin, 2015; Tucker et al., 2018). As such, researchers in diverse fields of science require methods allowing non-invasive measurement of movement, in different environments, across many animals simultaneously, and with high precision and accuracy (Dell et al., 2014).

The field of computer vision now allows researchers to automatically track subjects in a variety of contexts (Dell et al., 2014). Such automated tracking techniques drastically reduce the time and effort required to extract image-based data, allowing researchers to generate large and highly detailed datasets of animal movement and behaviour (e.g. Mersch, Crespi, & Keller, 2013; Attanasi et al., 2014). Methods for automated tracking abound in the computer vision literature, yet researchers often lack the know-how to implement these solutions for their specific tracking needs. Without a programming background, implementing such techniques can be daunting, and manual coding via direct observation often remains a preferred alternative, particularly for small-scale projects (Gomez-Marin, Paton, Kampff, Costa, & Mainen, 2014). However, while direct observation can produce informative data, it is often time-consuming, subjective, and challenging to replicate. Various proprietary software packages have been developed to provide simple automated tracking options (e.g. Noldus, Spink, & Tegenlenbosch, 2013), but these are expensive and often out of reach for many research groups. Clearly, there is a need for automated tracking solutions that are free, user-friendly and reliably provide image-based data acquisition solutions to record movement and behaviour.

A variety of open access tracking software already exist, designed to provide solutions for generic (e.g. Pérez-Escudero, Vicente-Page, Hinz, Arganda, & Polavieja, 2014; Rodriguez et al., 2018) and specific tracking problems (e.g. Branson, Robie, Bender, Perona, & Dickinson, 2009; Hewitt et al., 2018). However, automated tracking is achieved via different approaches, each one with its own strengths and weaknesses. Here, we propose Tracktor, a simple, open-source tracking software that can track multiple individuals simultaneously, single individuals in noisy environments, and produces a range of plots and summary statistics. Below, we detail its functioning, give examples of applications, and discuss its strengths and limitations. We also provide a detailed section on optimising video acquisition in the supplementary material.

## 2. Software

Tracktor is a python-based program that uses the OpenCV library (https://opencv.org/) for image processing. Briefly, Tracktor functions as a three-step process (Fig. 1): first, it automatically detects objects in videos through adaptive thresholding, and isolates the animals from other objects based on their size; second, it separates individuals from each other using k-means clustering; third, it maintains individual identities between frames using the Hungarian algorithm (Kuhn, 1955). A detailed description of the software is provided in the electronic supplementary material (ESM). There is no GUI (Graphical User Interface); all the commands are directly sent from scripts. However, all relevant input parameters are clearly explained and grouped in a designated cell at the start of the code, making Tracktor easy to use (Fig. S1). Additionally, the code is separated into sections and thoroughly annotated such that it is clear and accessible, even for an audience with minimal or no coding skills. For example, users with basic knowledge of the R statistical language will readily be able to use Tracktor. A detailed, step-by-step manual guiding users from installation to implementation is available on the Git repository: https://github.com/vivekhsridhar/tracktor.

**Figure 1.**
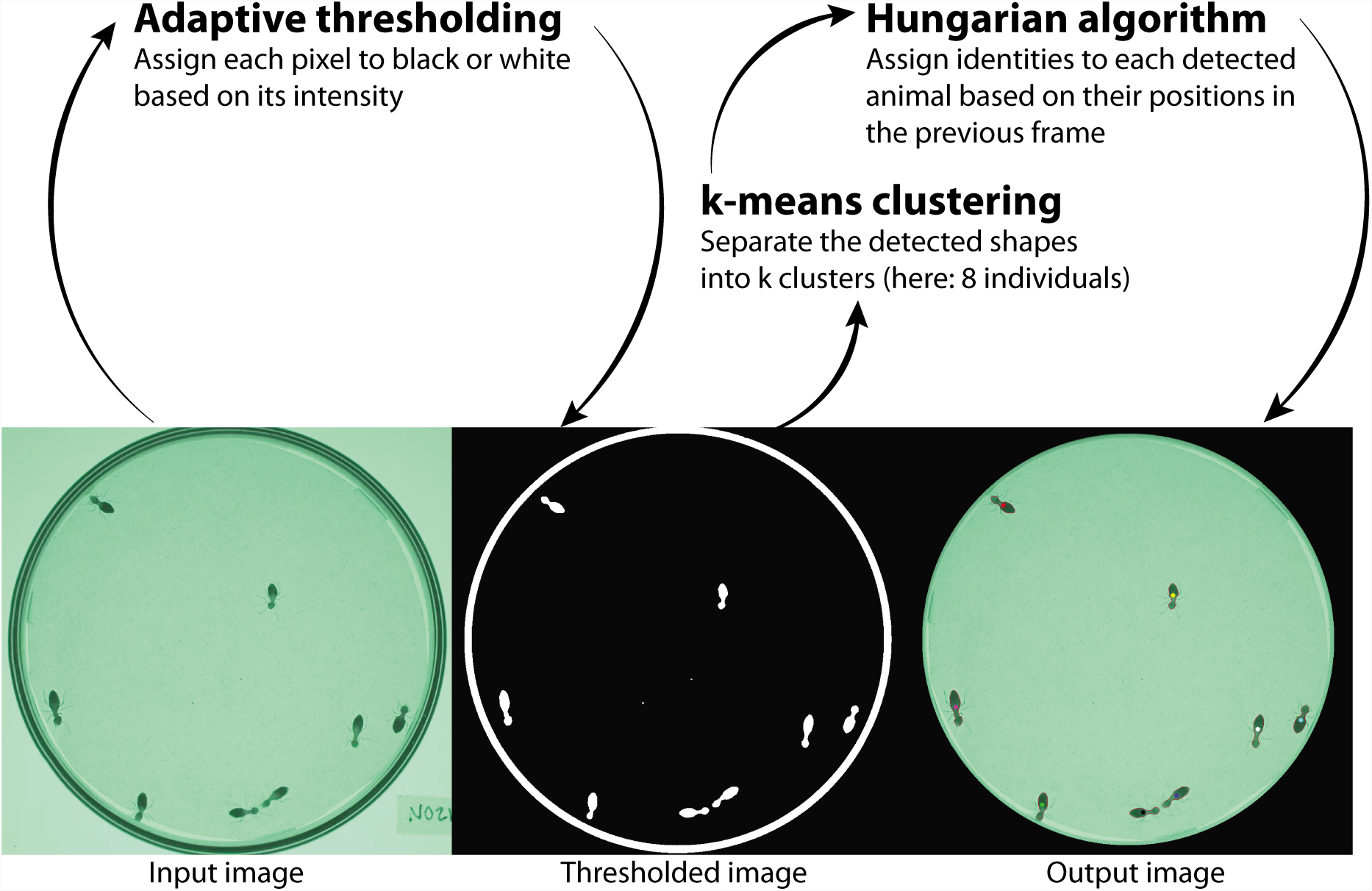
Overview of the different stages of the tracking procedure implemented in Tracktor, illustrated here for tracking 8 termites in a petri dish (example 3.4). First, adaptive thresholding is used to binarise the input image. A size rule is used to identify objects of interest (here, the largest object, the petri dish, is eliminated based on the size rule). Second, k-means clustering is used to separate objects if they are merged in the thresholded image. Third, individual identities are assigned using the Hungarian algorithm. Some steps are skipped depending on the tracking problem (e.g. k-means clustering is not implemented in examples 3.1, 3.2 and 3.3). See the ESM for a more detailed description of the software design.

## 3. Examples of applications

We provide four examples of common animal tracking problems that have successfully been solved with Tracktor. Each example is fully replicable and comprises annotated code to track the animals and produce relevant graphs and summary statistics. These examples can serve as the basis for solving similar tracking problems and can be readily adjusted to suit other problems with minimal modifications.

### 3.1 Fish fast-start escape response (1 individual)

Fast-start (i.e. burst) swimming performance in fishes is typically assessed by filming their escape behaviour in response to a mechano-acoustic stimulus falling inside the experimental tank (Domenici & Blake, 1997). When the stimulus hits the water surface, waves are created that propagate across the tank (Fig. 2a). These disturbances are problematic for automated tracking because water movements can be mistakenly detected as the animal, and several traditional image processing techniques (e.g. background subtraction) cannot be used in these conditions. As a result, most researchers code videos manually (e.g. Gingins, Roche, & Bshary, 2017; Shi et al., 2017), which is time consuming and prone to measurement error. Using a high speed (1000 fps) video of an escape response by the cleaning goby *Elacatinus prochilos*, we show that Tracktor can differentiate between animal movements and those of other shapes detected as potential objects. We provide summary statistics that are specific to fast-start analyses, and implement the Savitzky-Golay filter (Savitzky & Golay, 1964) to smooth noise and avoid erroneous measurements (Fig. 3a-c).

**Figure 2.**
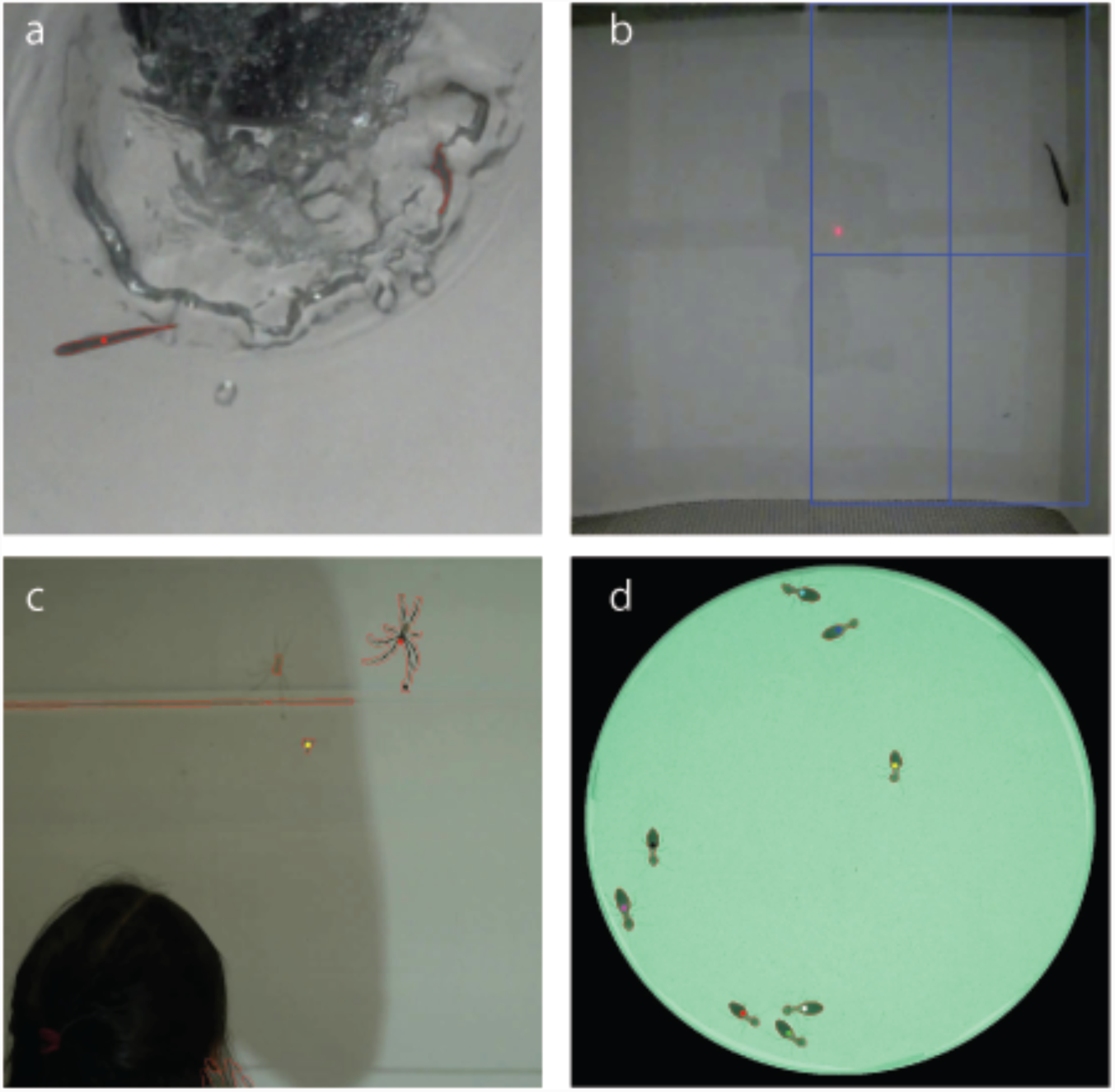
Examples of four different tracking problems solved with Tracktor: (a) a fast-start escape response by a single fish in an environment disturbed by a falling stimulus (example 3.1); (b) a single fish in a two-choice flume experiment with the region of interest depicted as a blue rectangle (example 3.2); (c) two spiders engaging in courtship in an environment disturbed by the experimenter (example 3.3); (d) collective behaviour of eight termites in an undisturbed environment (example 3.4).

**Figure 3.**
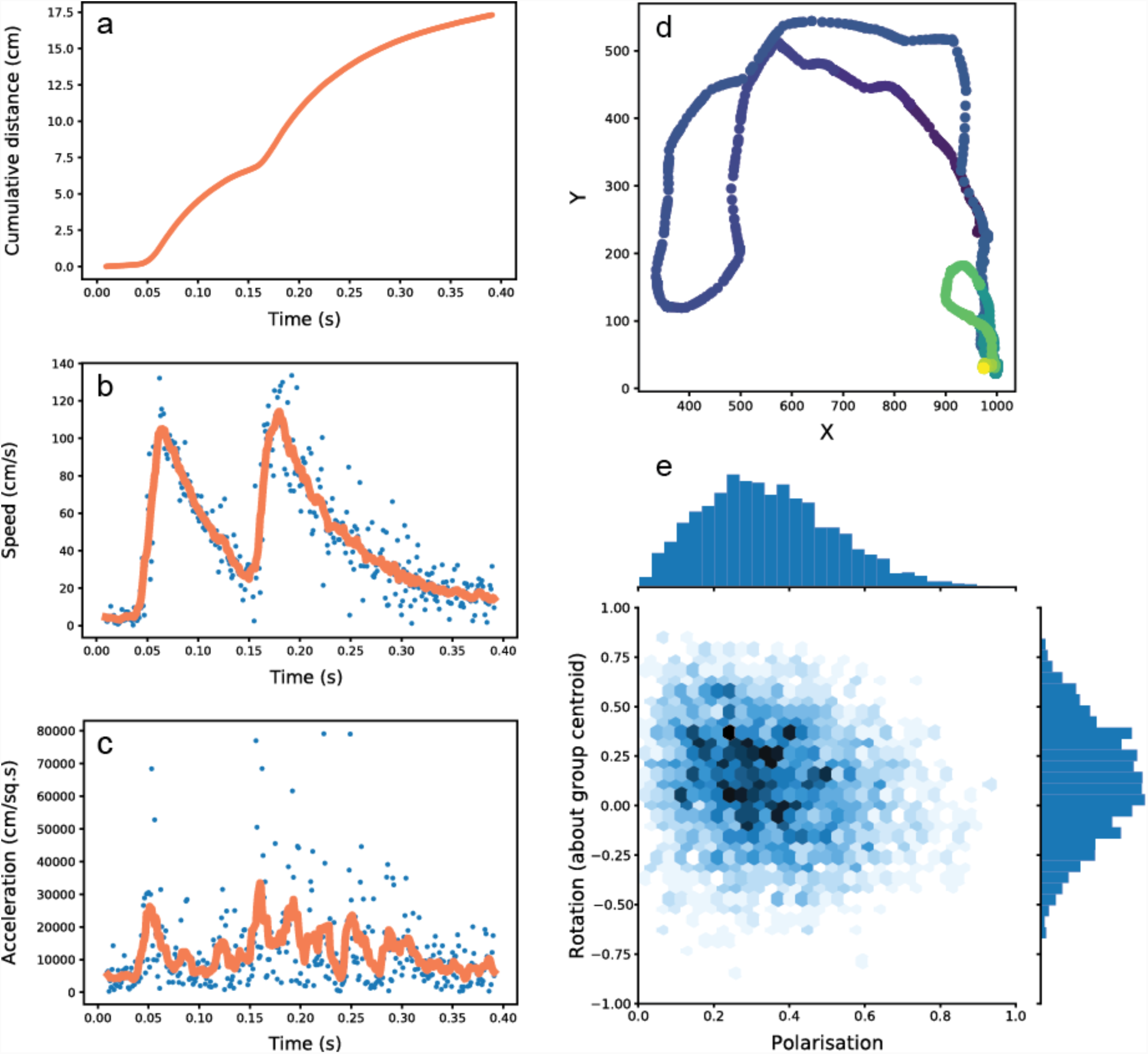
Example plots of summary statistics outputted by Tracktor. For the fish fast-start escape experiment (example 3.1), panels illustrate: **a)** cumulative distance covered [cm] vs. time [s], **b)** speed [cm s^-1^] vs. time [s], and **c)** acceleration [cm s^-2^] vs. time [s]. Blue dots are the raw data; orange lines represent the smoothed values of speed and acceleration using the Savitzky-Golay filter. Panel **d)** is the track [XY positions in pixels] for the flume choice experiment (example 3.2). Panel **e)** shows rotation vs. polarisation to assess the group’s dynamical state in the termite collective behaviour experiment.

### 3.2 Two-choice flume experiment (1 individual)

Researchers in behavioural ecology and ecotoxicology are often interested in measuring an animal’s avoidance or attraction to a given stimulus (e.g. olfactory or visual cues). Such choice tests are generally carried out by placing the test subject in a central arena with various choice options on different sides. The time spent in the vicinity of each option is then used as a proxy to measure preference or avoidance. Several experiments are based on this approach, for example to evaluate mate choice, kin and individual recognition, social tendencies, numerical abilities, and reaction to chemical cues (e.g. Hurst et al., 2001; Gómez-Laplaza & Gerlai, 2011; Fischer & Frommen, 2013; Jutfelt, Sundin, Raby, Krång, & Clark, 2017; Macario, Croft, Endler, & Darden, 2017). Here, we use Tracktor to track the movements of a single cleaning goby (*Elacatinus prochilos*) in a two-current choice flume with water containing the smells of different habitats. We first track the movements of the fish throughout the video. Then, we select an area by drawing a rectangle on the image, and measure the proportion of time the fish spent in the designated area (Fig. 2b).

### 3.3 Spider courtship behaviour (2 individuals)

Researchers often place two or more individuals in the same arena and record dyadic interactions, such as approach and avoidance behaviours. Such tracking problems are common in mate choice experiments, aggression, and sociability tests (Jolles, Boogert, Sridhar, Couzin, & Manica, 2017; Macario et al., 2017). Here we simultaneously track a male and a female spider (*Nephila senegalensis*) during courtship and provide distance between the two. In this example, Tracktor simultaneously tracks individuals of very different sizes (male *Nephila* are much smaller than females) and is robust to disturbances in the system. The key to successfully tracking objects of different sizes simultaneously is to ensure that the recording area is uncluttered since size thresholding cannot be used to its fullest extent when non-target objects of similar sizes are present in the arena (see ESM). Towards the end of the video sequence, the experimenter enters the field of view, casting a shadow on the setup. This does not affect tracking performance because Tracktor uses adaptive thresholding to cope with changes in lighting while disregarding the experimenter based on a specified size rule (see ESM).

### 3.4 Termite collective behaviour (8 individuals)

Studying collective behaviour requires recording the movements of multiple individuals simultaneously, resulting in large datasets. Automated tracking is extremely useful in this context, and many researchers already use this approach for their research by designing their own custom tracking solutions (Tunstrøm et al., 2013; Jolles et al., 2017). Here, we show that Tracktor can simultaneously track 8 termites (*Constrictotermes cyphergaster*) in a petri dish while maintaining individual identities. The challenge in this example is to correctly identify individuals: when two termites perform trophallaxis, thresholding results in the merging of their shapes. The k-means clustering technique implemented in Tracktor (see ESM) allows separating contours into a fixed number of clusters corresponding to the number of individuals in the video. Summary statistics are provided as standard measures of collective states such as polarisation and rotation of the group (Fig. 2e; see Couzin, Krause, James, Ruxton, & Franks, 2002; TunstrØm et al., 2013).

## 4. Strengths & limitations

Tracktor aims to provide an automated tracking tool that can be applied to many animals without the use of tags. Similar types of software (e.g. idTracker, Pérez-Escudero et al., 2014; BioTracker, Mönck et al., 2018; ToxTrac, Rodriguez et al., 2018) already exist, and we do not claim that Tracktor outperforms other software in all conditions. Rather, we emphasise that tracking problems and solutions are numerous and varied; hence, Tracktor provides an alternative that can fill the needs of many researchers when other software are not adequate. For instance, to our knowledge, Tracktor is the first tool to provide a robust and complete automated analysis of a fish fast-start escape response.

One of the common computer vision techniques to detect objects in videos is background subtraction, which relies on the assumption that the background is static while the animals to be tracked are mobile. Background subtraction has the advantage that it can deal with complex backgrounds and uneven illumination. However, if the animals remain static for a significant amount of time, detection will fail. For example, one of the eight termites (example 3.4) remained immobile throughout the recording. Additionally, it is not possible to use this technique when the background itself is moving, such as when measuring escape responses in fish (example 3.1). Under such conditions, Tracktor is thus likely to perform better than software using background subtraction (e.g. BioTracker).

We recognize that Tracktor’s lack of a GUI might initially discourage potential users from using the software. However, the growing prevalence of basic scripting/programming skills among evolutionary ecologists (e.g. in R) suggests that the absence of a GUI will not constitute a major impediment to most interested users, particularly given the simplicity of Tracktor’s code and the detailed user manual we provide alongside the software. A (python) code-based program also offers the additional advantage that it can easily be modified and improved by the community and run on all operating systems, which is often not the case for software with a GUI (e.g. ToxTrac).

Like many other software packages, Tracktor will not perform well if there are occlusions, or if animals go in and out of the camera’s field of view. Solving such problems requires algorithms that maintain identity by recognising individual features (e.g. idTracker) or assign IDs to tracklets when tracks are broken (e.g. ToxTrac). Such solutions require more computing power and result in longer processing times. Therefore, they were not implemented in the design of Tracktor.

## 5 Conclusion

Tracktor is a simple, rapid and versatile solution for many commonly encountered image-based tracking problems in animal behaviour. The software is robust to disturbances and allows users to track one or many individuals under a range of experimental conditions. Automated image-based tracking solutions such as Tracktor can greatly improve the study of animal movement and behaviour by reducing video processing time, avoiding observer bias, and allowing transparent, reproducible workflows.

## Acknowledgements

We thank I. D. Couzin and R. Bshary for supporting the development of Tracktor, M. Lampe, S. F. Garza, H. H. dos Santos and R. Mazzei for providing the example videos, I. D. Couzin and J. S. McClung for comments on the manuscript, and many colleagues who tested the software and provided feedback.

## Authors’ contributions

All authors designed and planned the study and contributed in writing the manuscript. V.H.S. developed the software.

## Data accessibility

The data used here are available at https://github.com/vivekhsridhar/tracktor.

